# DREAM-GNN: Dual-route embedding-aware graph neural networks for drug repositioning

**DOI:** 10.1101/2025.07.07.663530

**Authors:** Yanlong Zhao, Yixiao Chen, Jiawen Du, Quan Sun, Ren Wang, Can Chen

## Abstract

Drug repositioning presents a compelling strategy to accelerate therapeutic development by uncovering new indications for existing compounds. However, current computational methods are often limited in their ability to integrate heterogeneous biomedical data and model the intricate, multi-scale relationships underlying drug-disease associations, while large-scale experimental validation remains prohibitively resource-intensive. Here we present DREAM-GNN (Dual-Route Embedding-Aware Model for Graph Neural Networks), a multi-view deep graph learning framework that incorporates biomedical domain knowledge with two complementary graphs capturing both topological structure and feature similarity to enable accurate and biologically meaningful prediction of drug-disease associations. Extensive experiments on benchmark datasets demonstrate that DREAM-GNN significantly outperforms current state-of-the-art methods in recovering artificially removed repositioning candidates. These results establish DREAM-GNN as a robust and generalizable computational framework with broad potential to streamline drug discovery and advance precision medicine.

## Introduction

Drug repositioning, also known as drug repurposing, is the process of identifying new therapeutic uses for existing or previously failed drugs^1–6^. In contrast to traditional drug discovery, which is time-consuming, costly, and associated with a high failure rate, drug repositioning leverages established pharmacological and safety profiles to significantly reduce development time and expense required to bring treatments to market^7–9^. This paradigm has gained increasing attention in recent years as a valuable strategy for addressing unmet medical needs, particularly in complex or rare diseases where conventional discovery pipelines face limitations^10–13^. Notable successes, such as the repositioning of sildenafil, initially developed for angina and later approved for erectile dysfunction and pulmonary hypertension, highlight the clinical and commercial potential of this approach^14–16^. Recent applications extend to oncology, drug resistance mitigation, and personalized therapeutics, further reinforcing the promise of repositioning to diversify and modernize treatment landscapes^17–21^. Therefore, scalable, accurate, and data-driven methods are essential for drug repositioning, as large-scale experimental validation of candidate drug-disease associations remains prohibitively resource-intensive.

Computational methods have become central to modern repositioning efforts, enabling systematic prioritization of candidate associations across vast biomedical datasets. Early efforts predominantly relied on matrix factorization techniques, which project observed drug-disease interactions into a low-rank latent space while incorporating biological priors through drug and disease similarity matrices. Representative examples include DRRS^22^, which applies singular value thresholding to integrate disease-disease, drug-drug and drug-disease relationships, and BNNR^23^, which uses bounded nuclear norm regularization to fuse multiple biological similarity measures. Other variants, such as iDrug^24^ and SCMFDD^25^, exploit cross-network structures and semantic similarity constraints to improve biological interpretability. More recent extensions incorporate multi-view learning or genomic topology to enhance predictive performance^26, 27^. Yet, matrix factorization-based methods struggle to capture complex, nonlinear drug-disease interactions and become computationally demanding on large heterogeneous datasets.

The rapid proliferation of high-throughput biomedical data has fueled growing interest in machine learning approaches for drug repositioning^28–32^. These methods typically frame repositioning as a supervised learning task, predicting drug-disease associations from structured features derived from molecular, phenotypic, or network data. Pioneering models include PREDICT^33^, which integrates drug-drug and disease-disease similarity measures within a logistic regression framework to infer associations, alongside other methods that employ classifiers such as random forests to predict drug-target interactions^34^. These efforts laid the groundwork for more sophisticated deep learning models. For example, deepDR^35^ employs multi-modal autoencoders to learn high-level feature representations from heterogeneous drug-disease networks; CBPred^36^ leverages multi-path similarity to improve predictive accuracy; and DFDRNN^37^ introduces a deep fusion network that integrates diverse biological data sources through hierarchical deep learning modules. Despite these advances, such machine learning approaches generally treat neighboring nodes as independent, limiting their ability to model the complex relational patterns that underlie pharmacological and pathological processes.

Graph neural networks (GNNs) have recently emerged as leading approaches for drug repositioning by explicitly modeling both node features and network topology. Initial efforts focus on capturing local structural information and separately integrating similarity networks^38^, while more advanced architectures incorporate attention mechanisms and heterogeneous information fusion, exemplified by models such as LAGCN^39^ and DRHGCN^40^. Subsequent methods like DRWBNCF^41^ and PSGCN^42^ improve the modeling of context-specific dependencies through weighted convolution operations and subgraph-based classification strategies. The latest state-of-the-art frameworks, including DRAGNN^43^, AdaDR^44^, and DFDRNN^45^, incorporate weighted aggregation schemes, adaptive graph convolutions, and self-attention mechanisms to more effectively capture the complexity of biological interactions and improve model robustness. However, these GNN-based approaches still face significant challenges in effectively integrating heterogeneous multi-view data. More critically, they often fall short of fully leveraging available domain knowledge, such as drug chemical structures and disease semantics, limiting their biological interpretability and predictive power. These highlight a pressing need for novel GNN-based frameworks that can seamlessly integrate multi-view data and embed rich biochemical knowledge to accurately predict drug-disease associations.

To overcome these limitations, we introduce DREAM-GNN (Dual-Route Embedding-Aware Model for Graph Neural Networks), a novel framework that integrates multi-modal, pre-trained embeddings of drugs and diseases with two complementary graph encoders–one modeling network topology and the other capturing feature similarity–to enable accurate and biologically meaningful prediction of drug-disease associations. Through comprehensive evaluations across three benchmark datasets (Gdataset, Cdataset, and LRSSL), DREAM-GNN consistently outperforms state-of-the-art methods in identifying artificially removed drug repositioning candidates. These results demonstrate that DREAM-GNN delivers a biologically grounded and highly effective solution for computational drug repositioning.

## Materials and methods

### Overview of DREAM-GNN

We frame drug repositioning as heterogeneous link prediction on a bipartite drug-disease graph. DREAM-GNN first generates initial drug and disease embeddings by leveraging pre-trained large language-based models: ChemBERTa, ESM-2, and BioBERT. These embeddings are then refined in parallel through a relation-aware topology encoder and a feature-aware similarity encoder. Finally, an additive-attention gate integrates the dual views into a unified embedding, which is passed through a multi-layer perceptron to generate predicted scores for drug-disease pairs. The complete workflow of DREAM-GNN is illustrated in Fig. 1.

**Fig. 1:**
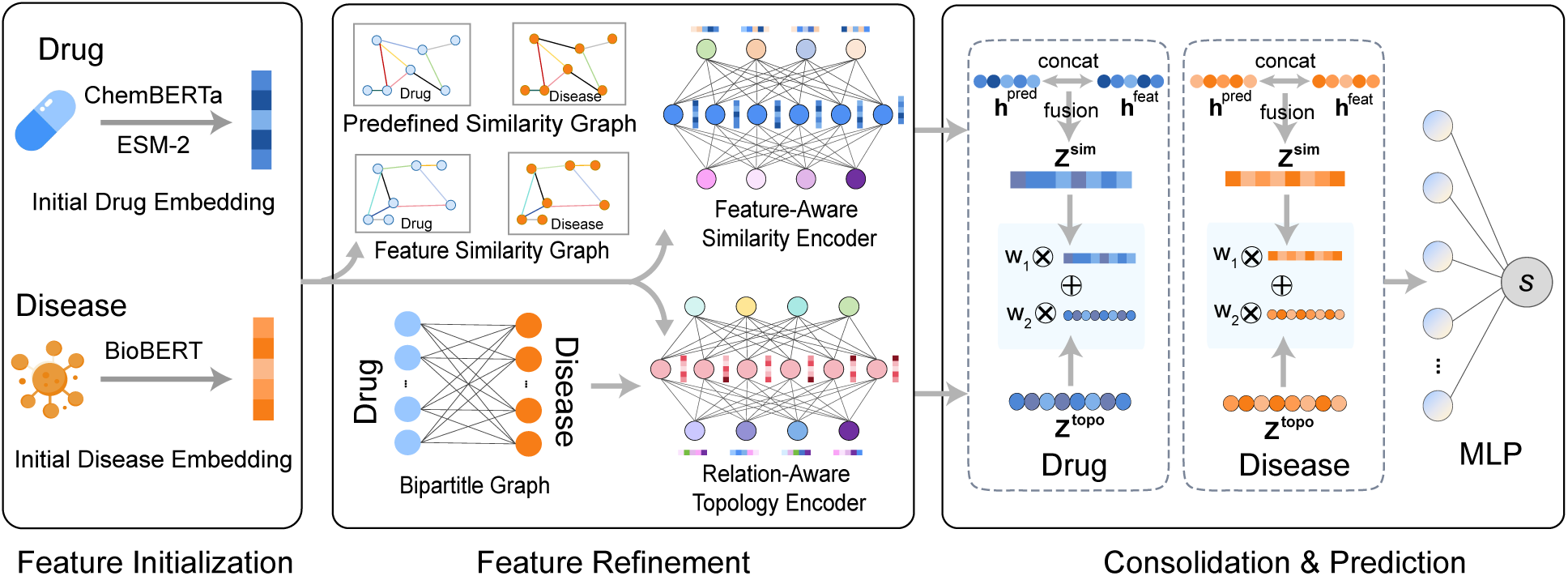
DREAM-GNN workflow. Feature Initialization: Drug embeddings are generated by encoding chemical SMILES with ChemBERTa and protein sequences with ESM-2. Disease embeddings are obtained using BioBERT on curated textual definitions. Feature Refinement: Two types of graphs are constructed, namely similarity graphs (domain-defined and embedding-derived *k*-nearest neighbor similarity) and a bipartite drug-disease interaction graph. A feature-aware similarity encoder applies GNNs separately to the drug and disease similarity graphs, while a relation-aware topology encoder refines node embeddings over the bipartite interaction graph. Consolidation & Prediction: Similarity-based and topology-informed embeddings are fused using a non-linear layer and attention mechanism, then decoded with an MLP to predict drug-disease associations.

### Feature initialization

Effective feature initialization is critical for capturing meaningful biological semantics and enabling accurate downstream prediction in graph-based drug repositioning models. To represent the diverse properties of drugs and diseases, DREAM-GNN employs modality-specific embedding strategies based on pre-trained large language models tailored to each entity type, including small-molecule drugs, protein therapeutics, and diseases. These heterogeneous embeddings are then harmonized into a unified feature space, ensuring compatibility and facilitating seamless integration within the graph neural network architecture.

### Drug initial embeddings

For small-molecule drugs, embeddings are derived from canonical SMILES (Simplified Molecular Input Line Entry System) strings using ChemBERTa^46^, a transformer model pre-trained on extensive chemical corpora. SMILES is a chemical notation system that represents molecular structures as compact ASCII text strings, encoding atoms, bonds, branching, and stereochemistry in a linear format that is particularly well-suited for computational processing and machine learning applications^47^. The canonical form ensures a unique, standardized representation for each molecule, eliminating structural redundancy and enabling consistent molecular encoding. Each canonical SMILES string is tokenized via byte-pair encoding, then truncated or padded to a fixed length of 510 tokens. These sequences are processed by ChemBERTa to generate contextual hidden states, which are pooled using masked mean pooling over non-padding tokens to produce a 1024-dimensional embedding. For protein therapeutics, a similar strategy is applied using amino acid sequences and ESM-2^48^, a 33-layer transformer protein language model. Sequences are augmented with beginning- and end-of-sequence tokens and fed into ESM-2, producing 1280-dimensional hidden states per token. Averaging the amino acid token states, excluding the boundary tokens, yields a 1280-dimensional embedding that captures the protein’s global semantic representation without positional bias.

To unify small-molecule and protein embeddings into a common feature space, the 1024-dimensional drug embeddings are zero-padded to match the 1280-dimensional protein vectors. These harmonized embeddings are then concatenated into a single feature matrix of size *N_d_*×1280, where each row corresponds to a drug, whether small molecule or protein therapeutic. To reduce redundancy and compress this unified matrix, principal component analysis is applied. Following common practice in dimensionality reduction, we select the top 768 components to balance computational efficiency with information retention, yielding a compact *N_d_* × 768 matrix that serves as the final drug embeddings for downstream graph-based inference.

### Disease initial embeddings

Each disease is encoded by converting its curated free-text description into a fixed-length embedding. Textual definitions from OMIM^49^ and MeSH^50^ are tokenized and input into the pre-trained BioBERT language model^51^, which generates a 768-dimensional contextual vector for each token, capturing nuanced biomedical semantics. A single representation per description is obtained by applying masked mean pooling to average only the vectors of substantive (non-padding) tokens, resulting in a 768-dimensional embedding for each disease. Repeating this process for all *N_m_* diseases produces a disease feature matrix with domain-aware embeddings ready for downstream analysis.

### Feature Refinement

DREAM-GNN refines initial drug and disease embeddings through a dual-route graph neural network that jointly captures two complementary sources of information: the explicit topology of known interactions and the implicit structure encoded in feature similarities.

### Relation-aware topology encoder

The relation-aware topology encoder operates on a bipartite drug-disease graph that incorporates two distinct edge types to distinguish evidence: ℛ_known_ represents clinically validated therapeutic associations, while ℛ_unknown_ denotes pairs with no known interaction, sampled as negative examples. A multi-layer GNN based on the graph convolutional matrix completion operator^52^ is employed to encode the relational topology. At each layer *l*, a node (drug or disease) embedding 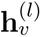 is refined by aggregating normalized messages from its neighbors under each relation type:

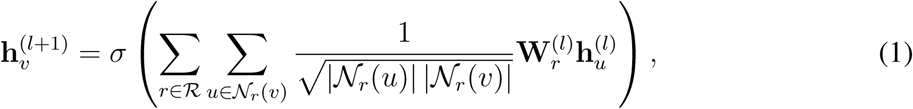

where 𝒩*_r_*(*v*) denotes the set of neighbors of node *v* under relation *r*, and *σ* is a nonlinear activation function. To balance expressiveness and parameter efficiency, each relation-specific weight matrix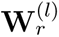 is factorized via basis decomposition: 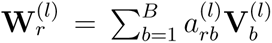, where 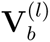 are shared basis matrices and 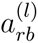 are relation-specific coefficients. The final topology-aware embedding 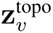 is obtained by averaging the node representations across all *L* layers: 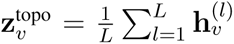, resulting in a multi-scale encoding that integrates signals from different neighborhood depths. This relation-aware topology encoder enables the model to distinguish between interaction types and effectively capture the structural heterogeneity of the drug-disease network.

### Feature-aware similarity encoder

The feature-aware similarity encoder is designed to capture rich, multi-granular similarity patterns that complement the explicit relational topology. For both drugs and diseases, two distinct *k*-nearest neighbor similarity graphs are constructed: (1) a pre-defined similarity graph, which for drugs is based on Tanimoto coefficients computed from chemical fingerprints, and for diseases, on semantic similarity scores derived from medical ontologies; and (2) a feature-based similarity graph, which encodes latent relationships by computing cosine similarity over the initial embeddings. Each of the four similarity graphs (two for drugs, two for diseases) is processed using a two-layer graph convolutional network. At each layer *l*, the node embeddings **H**^*(l)*^ are updated according to the propagation rule:

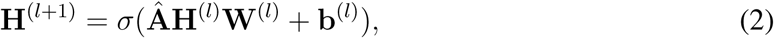

where 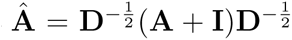 is the symmetrically normalized adjacency matrix with self-loops. For each entity type (drug or disease), the representations learned from the pre-defined similarity graph 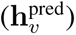 and the feature-derived graph 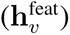 are fused to form a unified similarity embedding:

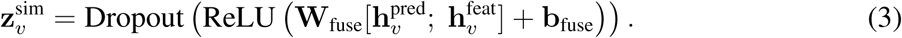

This feature-aware similarity encoder effectively captures both latent and pre-defined similarity patterns, enhancing the model’s capacity to represent complex and nuanced relationships within drug and disease entities.

### Consolidation and prediction

To obtain a unified representation for each drug and disease, the topology-aware embedding 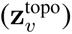 and similarity-aware embedding 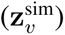 are combined using a learnable attention mechanism. The two embeddings are first stacked as 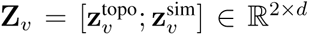. An attention network computes a normalized weight vector **w** = [*w*_1_ *w*_2_] ∈ ℝ^2^ that reflects the relative importance of each view:

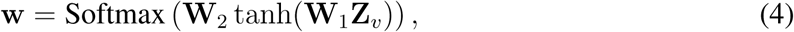

where **W**_1_ and **W**_2_ are learnable weight matrices. The final fused embedding is computed as a weighted sum of the two components:

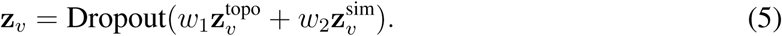

This attention-based fusion enables the model to adaptively integrate complementary structural and similarity information, yielding more informative and discriminative representations for both drugs and diseases. Finally, the fused embeddings of a drug *v_i_* (denoted by **z***_i_*) and a disease *v_j_* (denoted by **z***_j_*) are concatenated and passed through a three-layer multilayer perceptron (MLP) decoder, which outputs a single predicted score *s_ij_* = Sigmoid(MLP([**z***_i_*; **z***_j_*])). The model is trained end-to-end by minimizing the binary cross-entropy loss between the predicted scores and ground-truth labels, ensuring numerical stability and effective optimization.

### Advantages over existing methods

While rooted in the foundational design of GNNs, DREAM-GNN introduces targeted architectural and representational innovations that address the unique challenges of drug-disease association prediction. These advances enable it to surpass existing state-of-the-art methods in both accuracy and generalizability. First, DREAM-GNN emphasizes the integration of rich, multi-modal features derived from diverse sources (SMILES, protein sequences, disease texts) using specialized pre-trained models to create informative initial node embeddings. This contrasts with many prior methods that rely on simplistic node features such as one-hot encoding or random initialization. Second, DREAM-GNN adopts a dual-route architecture: a relation-aware topology encoder captures structural information from known drug-disease associations, while a feature-aware similarity encoder extracts relational patterns from similarity graphs constructed using both raw descriptors and learned embeddings. This design enables the model to fuse complementary insights from interaction topology and feature space geometry, an advantage over conventional single-graph approaches that often overlook such synergy. Furthermore, DREAM-GNN incorporates several robust learning strategies, including attention dropout, basis decomposition of weight matrices, and data augmentation via Gaussian feature perturbation and edge dropout, to enhance generalization and reduce the risk of overfitting.

## Experiments

### Datasets

To evaluate the performance of our proposed model, we utilize three widely used benchmark datasets: Gdataset^33^, Cdataset^53^ and LRSSL^54^. These datasets contain validated drug-disease associations and have been commonly adopted in drug repositioning studies.

- Gdataset^33^, often regarded as the gold-standard benchmark, contains 1,933 experimentally validated drug-disease pairs linking 593 small-molecule drugs from DrugBank to 313 Mendelian diseases cataloged in OMIM^55^. Its rigorous manual curation and balanced coverage of both drug and disease spaces have established it as a de facto standard for evaluating predictive models.
- Cdataset^53^, introduced by Luo et al., expands the evaluation scope to 2,352 drug-disease associations spanning 663 drugs and 409 diseases. Compared to Gdataset, it presents a denser interaction matrix and a slightly broader chemical space, allowing for a complementary assessment of model performance under more heterogeneous and noisier real-world conditions.
- The LRSSL dataset^54^ expands interaction coverage to 3,051 experimentally verified associations involving 763 FDA-approved drugs and 681 diseases. In addition to the bipartite drug-disease network, it includes three drug-drug similarity matrices based on chemical substructure, target protein domains, and Gene Ontology term overlap, as well as a semantic disease-disease similarity matrix derived from OMIM^55^, enabling methods that incorporate auxiliary similarity information.

### Baselines

To demonstrate the effectiveness of our proposed model, we compare its performance against several state-of-the-art methods for drug repositioning. These baseline methods represent a range of techniques, including matrix factorization- and GNN-based approaches.

- BNNR^23^ formulates drug repositioning as a matrix completion problem. It utilizes bounded nuclear norm regularization to predict potential drug-disease associations by recovering the underlying low-rank structure of the association matrix.
- DRAGNN^43^ is a recent GNN-based approach that improves drug repositioning predictions by integrating weighted local neighborhood information. It exploits the inherent graph structure of drug-disease interactions and assigns importance to neighboring nodes during message aggregation, enhancing the relevance of learned representations.
- AdaDR^44^ introduces an adaptive graph convolutional framework for drug repositioning, where the model dynamically learns or adjusts graph structures to better reflect the underlying topology of drug-disease interactions. This adaptability enables more effective information propagation tailored to the specific characteristics of the network.
- DFDRNN^37^ is a recent deep learning framework for drug-disease association prediction that introduces a deep fusion neural network to integrate diverse biological data sources. By combining heterogeneous information, the model aims to enhance predictive accuracy and uncover more reliable repositioning candidates.

### Experiment setup

#### Evaluation protocol

To rigorously assess the performance of our model and baseline methods, we adopt a 10-fold cross-validation strategy across all datasets. For each dataset, the set of known drug-disease associations (positive samples) and the complete set of unknown pairs (negative samples) are partitioned into 10 mutually exclusive folds. Unlike methods that employ downsampling strategies to balance class distributions, we utilize all available negative samples without subsampling, thus evaluating our model’s performance on the full scope of the drug-disease interaction space. This comprehensive approach better reflects real-world scenarios where the vast majority of drug-disease pairs are non-associations. In each cross-validation round, one fold is held out as the test set, while the remaining nine folds are used for training. Both positive and negative samples are split independently to ensure proper stratification. This process is repeated 10 times, ensuring that each fold is used exactly once for evaluation. Performance metrics are averaged across the folds to yield a robust estimate of generalization. To avoid information leakage, all graph structures and similarity matrices are constructed independently within each fold using only the training data available for that split.

#### Evaluation metrics

We assess the predictive performance using two standard metrics widely adopted in binary classification and drug repositioning tasks: the area under the receiver operating characteristic curve (AUROC) and the area under the precision-recall curve (AUPRC). AUROC evaluates the model’s ability to distinguish between positive and negative associations across various thresholds, while AUPRC is particularly informative for imbalanced datasets, measuring the trade-off between precision and recall. Higher values for both AUROC and AUPRC indicate better prediction performance.

#### Hyperparameters

We perform a grid search to identify optimal hyperparameter configurations using the LRSSL dataset. This strategic choice is motivated by two key considerations: (1) LRSSL is the largest and most comprehensive benchmark among the three datasets, containing 3,051 drug-disease associations across 763 drugs and 681 diseases, providing sufficient data diversity for robust parameter selection; (2) Hyperparameters optimized on this more complex and heterogeneous dataset are more likely to generalize well to the smaller and simpler Gdataset and Cdataset, following the principle that models tuned on challenging tasks typically transfer effectively to easier ones. Hyperparameter settings that yield the highest average AUPRC across cross-validation folds are selected for final evaluation. Table 1 summarizes the explored search space for each hyperparameter, with the optimal values (in bold) corresponding to those that achieve the best performance during training and validation on LRSSL. The identified optimal hyperparameters from LRSSL are then directly applied to Gdataset and Cdataset without further tuning, ensuring a fair comparison and demonstrating the generalizability of our approach.

**Table 1:** Hyperparameter selection for DREAM-GNN. Grid search was performed on the LRSSL dataset, with optimal values shown in bold.

| Component | Hyperparameter | Search Space |
| --- | --- | --- |
| Topology Encoder | Number of layers | {2, **3**, 4} |
|  | Aggregation units | {768, **1024**, 1536, 2048} |
|  | Output units | {96, **128**, 256} |
| | Basis decomposition ($B$) | {2, **4**, 8} |
| Similarity Encoder | Hidden units (layer 1) | {512, **768**, 1024} |
|  | Hidden units (layer 2) | {96, **128**, 256} |
| | $k$-NN neighbors ($k$) | {2, **4**, 8, 16} |
| Regularization | Dropout rate | {0.1, 0.2, **0.3**, 0.4} |
|  | Attention dropout | {0.0, **0.1**, 0.2} |
| Optimization | Learning rate | {1e-4, 1e-3, **2e-3**, 5e-3, 1e-2} |
|  | Weight decay | {1e-6, **1e-5**, 1e-4} |

### Performance Comparison and Analysis

DREAM-GNN signficantly outperforms existing state-of-the-art methods across all the three benchmark datasets, demonstrating both strong predictive accuracy and stability under varying data conditions (Fig. 2). On the widely used Cdataset, DREAM-GNN achieves median scores of 72.20% AUPRC and 97.38% AUROC, surpassing the second-best method DFDRNN by 6.07 percentage points in AUPRC (*P* -value = 3.2E-9, paired *t*-test) and 0.62 points in AUROC (*P* - value = 2.5E-8). Other baselines including AdaDR, DRAGNN, and BNNR lag considerably behind with poor predictive power. Performance gains are even more striking on the more heterogeneous Gdataset, where DREAM-GNN achieves 73.72% AUPRC and 98.27% AUROC. In comparison, DFDRNN trails by 20.86 points in AUPRC (*P* -value = 9.5E-13) and 2.90 points in AUROC (*P* - value = 3.7E-11), indicating DREAM-GNN’s capacity to generalize effectively across noisier and more complex interactions. Evaluation on the LRSSL dataset further underscores DREAM-GNN’s robustness. It reaches 55.24% AUPRC and 97.86% AUROC, outperforming DFDRNN by 12.44 points in AUPRC (*P* -value = 7.9E-11) and 1.83 points in AUROC (*P* -value = 3.2E-8).

**Fig. 2:**
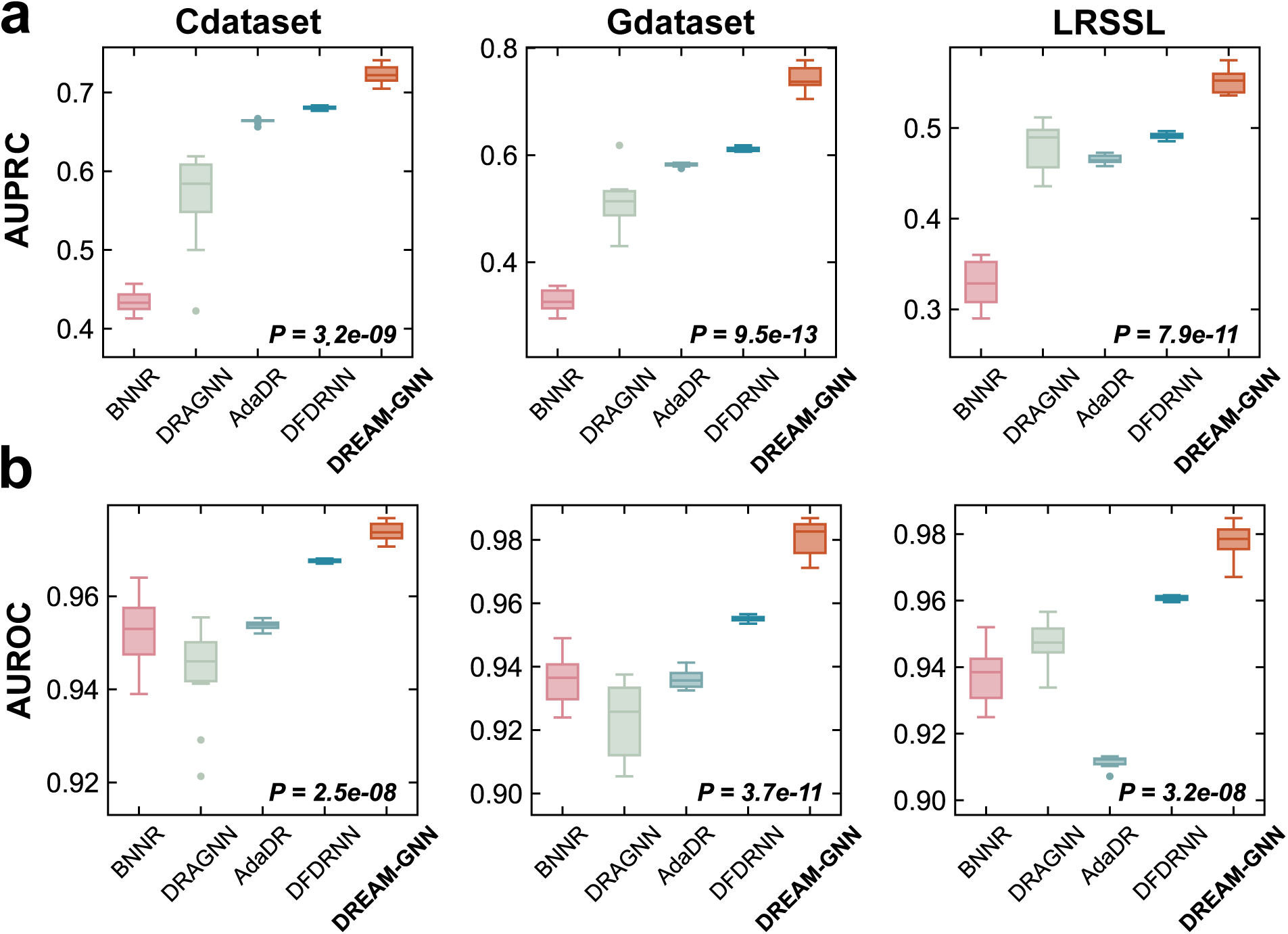
Comparison of DREAM-GNN with existing methods across three datasets. DREAM-GNN outperforms BNNR, DRAGNN, AdaDR, DFDRNN on both AUPRC (**a**) and AUROC (**b**) metrics across Cdataset, Gdataset, and LRSSL. Boxplots reflect performance distributions over 10-fold cross-validation. *P* -values indicate statistical significance from paired t-tests against the strongest baseline for each dataset.

Notably, DREAM-GNN delivers substantial and consistent gains in AUPRC across all evaluated datasets, underscoring its effectiveness in settings characterized by severe class imbalance, a common hallmark of drug-disease association tasks. Unlike AUROC, AUPRC is more sensitive to the performance on the minority (positive) class and thus better reflects real-world utility in identifying novel therapeutic associations. On the challenging Gdataset, for example, DREAM-GNN achieves an AUPRC of 73.72%, representing a 20.86-point improvement over the next-best method DFDRNN. Similar trends are observed on Cdataset and LRSSL, with DREAM-GNN outperforming all baselines by wide and statistically significant margins. Furthermore, DREAM-GNN exhibits significantly lower variability compared to methods like BNNR and DRAGNN, indicating superior generalizability and stability. These improvements suggest that the model effectively prioritizes biologically meaningful associations amidst overwhelming noise, crucial for practical drug repositioning efforts.

### Ablation Study

To systematically investigate the effectiveness and contributions of each component within the DREAM-GNN framework, we conduct a comprehensive ablation study on the LRSSL dataset (Table 2). Specifically, we examine the following four critical variants: (1) DREAM-GNN-w/o-attention-dropout, which removes the dropout regularization from the attention fusion mechanism; (2) DREAM-GNN-w/o-parameter-sharing, where parameters in the graph convolutional layers are independently learned without sharing; (3) DREAM-GNN-w/o-augmentation, omitting all data augmentation strategies, including Gaussian feature noise and edge dropout; and (4) DREAM-GNN-w/o-node-feature-embedding, which replaces the pretrained embeddings derived from ChemBERTa, ESM-2, and BioBERT with random initialization.

**Table 2:** Ablation study results for DREAM-GNN. Average test AUROC and AUPRC are reported with standard deviation over multiple runs.

| **Model / Strategy** | **AUROC** | **AUPRC** |
| --- | --- | --- |
| DREAM-GNN | $0.9843 \pm 0.0048$ | $0.5748 \pm 0.0568$ |
| w/o Attention Dropout | $0.9799 \pm 0.0043$ | $0.5644 \pm 0.0475$ |
| w/o Parameter Sharing | $0.9758 \pm 0.0053$ | $0.3546 \pm 0.1489$ |
| w/o Augmentation | $0.9716 \pm 0.0253$ | $0.5409 \pm 0.0882$ |
| w/o node feature embedding | $0.9132 \pm 0.0445$ | $0.4625 \pm 0.0118$ |

The results clearly illustrate that each component plays an essential role in the overall predictive performance of DREAM-GNN. Removing pre-trained node embeddings leads to the most pronounced performance drop, substantially decreasing both AUPRC and AUROC (AUROC: 0.9132 ± 0.0445, AUPRC: 0.4625 ± 0.0118), indicating the critical importance of incorporating rich, domain-specific feature information for capturing intrinsic biological semantics. Similarly, disabling parameter sharing markedly deteriorates AUPRC, underscoring the benefits of parameter efficiency and improved generalization achieved through shared convolutional weights. The absence of data augmentation strategies, such as feature noise and edge dropout, also negatively impacts model robustness, suggesting these augmentations effectively mitigate overfitting. Finally, removing dropout in the attention fusion mechanism leads to a moderate yet noticeable performance reduction, highlighting its role in enhancing the stability and generalization capabilities of the learned attention weights. Collectively, these findings confirm that each proposed component contributes meaningfully to its overall predictive accuracy, robustness, and generalization capability.

## Discussion

In this work, we present DREAM-GNN, a dual-route embedding-aware graph neural network framework tailored to tackle the challenges of data heterogeneity in computational drug repositioning. DREAM-GNN systematically integrates rich, modality-specific embeddings derived from state-of-the-art pre-trained language models, including ChemBERTa, ESM-2, and BioBERT, to establish a comprehensive initial representation of drugs and diseases. Its dual-encoder architecture jointly models explicit interaction topologies and implicit semantic similarities, while a learnable attention mechanism dynamically fuses these complementary views. This design enables DREAM-GNN to learn robust and discriminative representations for accurate prediction of novel drug-disease associations. Extensive evaluation across three large-scale benchmark datasets demonstrates that DREAM-GNN consistently outperforms existing methods, highlighting its effectiveness and generalizability.

By leveraging DREAM-GNN, biomedical researchers are equipped with a powerful and generalizable framework for uncovering novel drug-disease associations with high precision, even in the face of noisy, incomplete, or heterogeneous data, a common challenge in real-world biomedical research. The model’s integration of biochemical, genomic, and clinical knowledge through pre-trained embeddings enables the discovery of non-obvious therapeutic connections that may be overlooked by traditional approaches. This capability not only facilitates hypothesis generation for experimental validation but also supports the repurposing of approved or shelved compounds for emerging diseases, rare disorders, or personalized treatment strategies. In doing so, DREAM-GNN advances the translational utility of computational discovery, offering a scalable tool to bridge data-driven insights with therapeutic innovation.

While DREAM-GNN establishes a flexible and effective framework, there are areas for future enhancement. First, the model currently integrates features from chemical structures, protein sequences, and disease descriptions. Its predictive power could be further enhanced by incorporating additional biomedical knowledge, such as gene expression profiles, protein-protein interaction networks, and drug side-effect data, to construct a more comprehensive and informative heterogeneous graph. Second, although the attention mechanism offers some level of interpretability, future work should focus on developing more advanced techniques to translate the model’s predictions into biologically verifiable hypotheses regarding drug mechanisms of action. Third, as biomedical datasets continue to grow, exploring graph sampling techniques and memory optimization strategies will be crucial to ensure the framework remains computationally efficient when scaling to web-scale knowledge graphs. Finally, beyond computational validation, the ultimate utility of DREAM-GNN depends on its biological relevance. Therefore, future work should prioritize experimental and clinical validation of model predictions to confirm their practical value in real-world drug discovery and translational medicine settings.

## Code Availability

The source code for DREAM-GNN is publicly available at: https://github.com/Ryan-Yanlong/DREAM-GNN.

## References

1. Nathan C Baker, Sean Ekins, Antony J Williams, and Alexander Tropsha. A bibliometric review of drug repurposing. Drug Discovery Today, 23(3):661–672, 2018.

2. Sudeep Pushpakom, Francesco Iorio, Patrick A Eyers, K Jane Escott, Shirley Hopper, Andrew Wells, Andrew Doig, Tim Guilliams, Joanna Latimer, Christine McNamee, et al. Drug repurposing: progress, challenges and recommendations. Nature reviews Drug discovery, 18(1):41–58, 2019.

3. Michael J Barratt and Donald E Frail. Drug repositioning: Bringing new life to shelved assets and existing drugs. John Wiley & Sons, 2012.

4. Hanqing Xue, Jie Li, Haozhe Xie, and Yadong Wang. Review of drug repositioning approaches and resources. International journal of biological sciences, 14(10):1232, 2018.

5. Ali Ezzat, Min Wu, Xiao-Li Li, and Chee-Keong Kwoh. Computational prediction of drug–target interactions using chemogenomic approaches: an empirical survey. Briefings in bioinformatics, 20(4):1337–1357, 2019.

6. Tamer N Jarada, Jon G Rokne, and Reda Alhajj. A review of computational drug repositioning: strategies, approaches, opportunities, challenges, and directions. Journal of cheminformatics, 12:1–23, 2020.

7. Jiao Li, Si Zheng, Bin Chen, Atul J Butte, S Joshua Swamidass, and Zhiyong Lu. A survey of current trends in computational drug repositioning. Briefings in bioinformatics, 17(1):2–12, 2016.

8. Joseph A DiMasi. New drug development in the united states from 1963 to 1999. Clinical Pharmacology & Therapeutics, 69(5):286–296, 2001.

9. Allen Krantz. Diversification of the drug discovery process. Nature biotechnology, 16(13):1294–1294, 1998.

10. Charlotte Harrison. Coronavirus puts drug repurposing on the fast track. Nature biotechnology, 38(4):379–381, 2020.

11. Yadi Zhou, Fei Wang, Jian Tang, Ruth Nussinov, and Feixiong Cheng. Artificial intelligence in covid-19 drug repurposing. The Lancet Digital Health, 2(12):e667–e676, 2020.

12. Helen I Roessler, Nine VAM Knoers, Mieke M van Haelst, and Gijs van Haaften. Drug repurposing for rare diseases. Trends in pharmacological sciences, 42(4):255–267, 2021.

13. Divya Sardana, Cheng Zhu, Minlu Zhang, Ranga C Gudivada, Lun Yang, and Anil G Jegga. Drug repositioning for orphan diseases. Briefings in bioinformatics, 12(4):346–356, 2011.

14. Jean-Pierre Jourdan, Ronan Bureau, Christophe Rochais, and Patrick Dallemagne. Drug repositioning: a brief overview. Journal of Pharmacy and Pharmacology, 72(9):1145–1151, 09 2020.

15. Natalia Novac. Challenges and opportunities of drug repositioning. Trends in pharmacological sciences, 34(5):267–272, 2013.

16. Zhichao Liu, Hong Fang, Kelly Reagan, Xiaowei Xu, Donna L Mendrick, William Slikker Jr, and Weida Tong. In silico drug repositioning–what we need to know. Drug discovery today, 18(3-4):110–115, 2013.

17. Javier Setoain, Monica Franch, Marta Martínez, Daniel Tabas-Madrid, Carlos OS Sorzano, Annette Bakker, Eduardo Gonzalez-Couto, Juan Elvira, and Alberto Pascual-Montano. Nffinder: an online bioinformatics tool for searching similar transcriptomics experiments in the context of drug repositioning. Nucleic acids research, 43(W1):W193–W199, 2015.

18. Waleed Younis, Shankar Thangamani, and Mohamed N Seleem. Repurposing non-antimicrobial drugs and clinical molecules to treat bacterial infections. Current pharmaceutical design, 21(28):4106–4111, 2015.

19. Yvonne Y Li and Steven JM Jones. Drug repositioning for personalized medicine. Genome medicine, 4:1–14, 2012.

20. Roberto Wuerth, Stefano Thellung, Adriana Bajetto, Michele Mazzanti, Tullio Florio, and Federica Barbieri. Drug-repositioning opportunities for cancer therapy: novel molecular targets for known compounds. Drug discovery today, 21(1):190–199, 2016.

21. Guangxu Jin, Changhe Fu, Hong Zhao, Kemi Cui, Jenny Chang, and Stephen TC Wong. A novel method of transcriptional response analysis to facilitate drug repositioning for cancer therapy. Cancer research, 72(1):33–44, 2012.

22. Huimin Luo, Min Li, Shaokai Wang, Quan Liu, Yaohang Li, and Jianxin Wang. Computational drug repositioning using low-rank matrix approximation and randomized algorithms. Bioinformatics, 34(11):1904–1912, 2018.

23. Mengyun Yang, Huimin Luo, Yaohang Li, and Jianxin Wang. Drug repositioning based on bounded nuclear norm regularization. 35(14):i455–i463, 2019.

24. Huiyuan Chen, Feixiong Cheng, and Jing Li. idrug: Integration of drug repositioning and drug-target prediction via cross-network embedding. PLoS computational biology, 16(7):e1008040, 2020.

25. Wen Zhang, Xiang Yue, Weiran Lin, Wenjian Wu, Ruoqi Liu, Feng Huang, and Feng Liu. Predicting drug-disease associations by using similarity constrained matrix factorization. BMC bioinformatics, 19:1–12, 2018.

26. Yixin Yan, Mengyun Yang, Haochen Zhao, Guihua Duan, Xiaoqing Peng, and Jianxin Wang. Drug repositioning based on multi-view learning with matrix completion. Briefings in Bioinformatics, 23(3):bbac054, 2022.

27. Wen Dai, Xi Liu, Yibo Gao, Lin Chen, Jianglong Song, Di Chen, Kuo Gao, Yongshi Jiang, Yiping Yang, Jianxin Chen, et al. Matrix factorization-based prediction of novel drug indications by integrating genomic space. Computational and mathematical methods in medicine, 2015(1):275045, 2015.

28. Antonio Lavecchia and Carmen Cerchia. In silico methods to address polypharmacology: current status, applications and future perspectives. Drug Discovery Today, 21(2):288–298, 2016.

29. Maria Koromina, Maria-Theodora Pandi, and George P Patrinos. Rethinking drug repositioning and development with artificial intelligence, machine learning, and omics. Omics: a journal of integrative biology, 23(11):539–548, 2019.

30. Lijun Cai, Jiaxin Chu, Junlin Xu, Yajie Meng, Changcheng Lu, Xianfang Tang, Guanfang Wang, Geng Tian, and Jialiang Yang. Machine learning for drug repositioning: Recent advances and challenges. Current Research in Chemical Biology, 3:100042, 2023.

31. Francesco Napolitano, Yan Zhao, Vânia M Moreira, Roberto Tagliaferri, Juha Kere, Mauro D’Amato, and Dario Greco. Drug repositioning: a machine-learning approach through data integration. Journal of cheminformatics, 5:1–9, 2013.

32. Fei Wang, Yulian Ding, Xiujuan Lei, Bo Liao, and Fang-Xiang Wu. Machine learning and deep learning strategies in drug repositioning. Current Bioinformatics, 17(3):217–237, 2022.

33. Assaf Gottlieb, Gideon Y Stein, Eytan Ruppin, and Roded Sharan. Predict: a method for inferring novel drug indications with application to personalized medicine. Molecular Systems Biology, 7:496, 2011.

34. Dong-Sheng Cao, Liu-Xia Zhang, Gui-Shan Tan, Zheng Xiang, Wen-Bin Zeng, Qing-Song Xu, and Alex F Chen. Computational prediction of drug target interactions using chemical, biological, and network features. Molecular informatics, 33(10):669–681, 2014.

35. Xiangxiang Zeng, Siyi Zhu, Xiangrong Liu, Yadi Zhou, Ruth Nussinov, and Feixiong Cheng. deepdr: a network-based deep learning approach to in silico drug repositioning. Bioinformatics, 35(24):5191–5198, 2019.

36. Ping Xuan, Yilin Ye, Tiangang Zhang, Lianfeng Zhao, and Chang Sun. Convolutional neural network and bidirectional long short-term memory-based method for predicting drug–disease associations. Cells, 8(7):705, 2019.

37. Enqiang Zhu, Xiang Li, Chanjuan Liu, and Nikhil R. Pal. Boosting drug-disease association prediction for drug repositioning via dual-feature extraction and cross-dual-domain decoding, 2025.

38. Jin Li, Sai Zhang, Tao Liu, Chenxi Ning, Zhuoxuan Zhang, and Wei Zhou. Neural inductive matrix completion with graph convolutional networks for mirna-disease association prediction. Bioinformatics, 36(8):2538–2546, 2020.

39. Zhouxin Yu, Feng Huang, Xiaohan Zhao, Wenjie Xiao, and Wen Zhang. Predicting drug– disease associations through layer attention graph convolutional network. Briefings in bioinformatics, 22(4):bbaa243, 2021.

40. Lijun Cai, Changcheng Lu, Junlin Xu, Yajie Meng, Peng Wang, Xiangzheng Fu, Xiangxiang Zeng, and Yansen Su. Drug repositioning based on the heterogeneous information fusion graph convolutional network. Briefings in bioinformatics, 22(6):bbab319, 2021.

41. Yajie Meng, Changcheng Lu, Min Jin, Junlin Xu, Xiangxiang Zeng, and Jialiang Yang. A weighted bilinear neural collaborative filtering approach for drug repositioning. Briefings in bioinformatics, 23(2):bbab581, 2022.

42. Xinliang Sun, Bei Wang, Jie Zhang, and Min Li. Partner-specific drug repositioning approach based on graph convolutional network. IEEE Journal of Biomedical and Health Informatics, 26(11):5757–5765, 2022.

43. Yajie Meng, Yi Wang, Junlin Xu, Changcheng Lu, Xianfang Tang, Tao Peng, Bengong Zhang, Geng Tian, and Jialiang Yang. Drug repositioning based on weighted local information augmented graph neural network. Briefings in Bioinformatics, 25(1):bbad431, 2024.

44. Xinliang Sun, Xiao Jia, Zhangli Lu, Jing Tang, and Min Li. Drug repositioning with adaptive graph convolutional networks. Bioinformatics, 40(1):btad748, 2024.

45. Enqiang Zhu, Xiang Li, Chanjuan Liu, and Nikhil R Pal. Dfdrnn: A dual-feature based neural network for drug repositioning. arXiv e-prints, pages arXiv–2407, 2024.

46. B. Chithra, A. G. Prakash, T. Emmanuel, et al. Chemberta: Training and interpreting large language models for chemical text. arXiv preprint arXiv:2010.09885, 2020.

47. David Weininger. Smiles, a chemical language and information system. 1. introduction to methodology and encoding rules. Journal of Chemical Information and Computer Sciences, 28(1):31–36, 1988.

48. Zeming Lin, Halil Akin, Roshan Rao, Brian Hie, Zhongkai Zhu, Wenting Lu, Nikita Smetanin, Allan dos Santos Costa, Maryam Fazel-Zarandi, Tom Sercu, Sal Candido, et al. Language models of protein sequences at the scale of evolution enable accurate structure prediction. bioRxiv, 2022.

49. Ada Hamosh, Alan F Scott, Joanna S Amberger, Catharine A Bocchini, and Victor A McKusick. Online mendelian inheritance in man (omim), a knowledgebase of human genes and genetic disorders. Nucleic Acids Research, 33(suppl 1):D514–D517, 2005.

50. Carolyn E Lipscomb. Medical subject headings (mesh). Bulletin of the Medical Library Association, 88(3):265–266, 2000.

51. Jinhyuk Lee, Wonjin Yoon, Sungdong Kim, Donghyeon Kim, Sunkyu Kim, Chan Ho So, and Jaewoo Kang. BioBERT: a pre-trained biomedical language representation model for biomedical text mining. Bioinformatics, 36(4):1234–1240, 2020.

52. Rianne van den Berg, Thomas N. Kipf, and Max Welling. Graph convolutional matrix completion, 2017.

53. Hongwu Luo, Jing Wang, Mingyang Li, Jin Luo, Pei Ni, Kan Zhao, Fangxiang Wu, Yi Pan, and Qinghua Jiang. Drug repositioning based on comprehensive similarity measures and bi-random walk algorithm. Bioinformatics, 32(17):2664–2671, 2016.

54. Xujun Liang, Pengfei Zhang, Lu Yan, Ying Fu, Fang Peng, Lingzhi Qu, Meiying Shao, Yongheng Chen, and Zhuchu Chen. Lrssl: predict and interpret drug–disease associations based on data integration using sparse subspace learning. Bioinformatics, 33(8):1187–1196, 2017.

55. Judith S. Amberger, Carol A. Bocchini, Alan F. Scott, and Ada Hamosh. OMIM.org: Leveraging knowledge across phenotype–gene relationships. Nucleic Acids Research, 47(D1):D1038–D1043, 2019.

